# L-serine diet restores impaired adult neurogenesis in the hippocampus of 3xTg-AD mice

**DOI:** 10.64898/2026.04.27.721014

**Authors:** Emmanuel Than-Trong, Lucille Torres, Mylène Gaudin-Guérif, Caroline Jan, Aurélie Ghettas, Aurélie Amadio, The Brainbank Neuro-CEB Neuropathology Network, Stéphane H.R. Oliet, Aude Panatier, Gilles Bonvento

**Author notes:** **Corresponding author:** Gilles Bonvento, PhD, Institut des Neurosciences Paris-Saclay (NeuroPSI) France.

## Abstract

Altered adult neurogenesis is reported in Alzheimer’s disease (AD) in humans and rodent models, though the mechanisms remain unclear. L-serine, a non-essential amino acid that plays a critical role in cell proliferation and survival, is produced by neuroepithelial cells and radial glia in the developing brain, as well as by astrocytes and neural precursors in the adult brain. Its production is altered in AD, particularly in the hippocampus. We sought to determine whether a deficiency of L-serine availability contributes to the reduced adult neurogenesis in AD. We confirm that phosphoglycerate dehydrogenase (PHGDH), the rate-limiting enzyme of the L-serine biosynthetic pathway, is expressed by neural stem cells (NSCs) of the mouse dentate gyrus (DG). We further report PHGDH expression in cells with somata located in the subgranular zone (SGZ) of the human DG and displaying the typical radial morphology associated with NSCs in rodents. We observed a significant decrease in the number of proliferating cells (proliferating cell nuclear antigen, PCNA+) as well as immature neurons (doublecortin, DCX+) in the DG of 12-month-old 3xTg-AD mice compared to their age-matched controls. Importantly, chronic dietary supplementation with a L-serine-enriched diet for 8 months significantly increased plasma L- and D-serine levels and partially rescued adult neurogenesis deficits in 3xTg-AD mice, while having no significant impact on the progression of amyloidosis. Our results suggest that chronic metabolic impairment in L-serine production, and the resulting shortage of D-serine, likely contributes to reduced survival of newborn neurons in the DG of 3xTg-AD mice.

## 1 Introduction

Alzheimer’s disease (AD) is a progressive neurodegenerative disorder that impairs memory and cognitive function, gradually disrupting daily life. AD features two primary pathological hallmarks: extracellular plaques, mainly composed of amyloid-beta 42, and intracellular neurofibrillary tangles (NFTs) formed from hyperphosphorylated TAU ^1^. Functional and structural disruption of synapses is central to the disease process ^2, 3^, particularly in the hippocampus, one of the first brain regions to be affected by AD pathology ^4^. The dentate gyrus (DG), a hippocampal subregion implicated in learning and memory, notably in pattern separation, contains the “neurogenic niche,” a specialized environment where neural stem cells (NSCs) continue to produce new neurons in the adult brain of most mammalian species studied to date ^5-7^, including humans ^8, 9.^ This process is known as adult hippocampal neurogenesis (AHN). While the extent and functional significance of AHN in humans remain a matter of debate ^10^, several recent studies have reported a reduction in the number of immature neurons in the DG of AD patients ^11-14^. Similarly, rodent models of AD demonstrate a decline in AHN preceding the appearance of the classical pathological hallmarks of the disease ^15^. The precise molecular mechanisms underlying the impairment of AHN in AD remain unclear, but are likely linked to a generalized disruption of DG homeostasis ^16^.

Hippocampal neurogenesis is tightly regulated by niche signals, with energy metabolism directing NSC proliferation and maturation into fully differentiated neurons ^17^. The metabolic processes underlying AHN are governed not only by local intracellular energy metabolism ^18^ but also by systemic energy availability and the organism’s integrated physiological responses to energy status.

Disrupted energy metabolism, observed in many pathologies including neurodegenerative diseases, may impair NSC function, and is altered early in AD ^19, 20.^ We have previously demonstrated that the glycolytic flux is impaired in the hippocampus of young 3xTg-AD mice, leading to reduced astrocytic production of L-serine, an amino acid generated through the phosphorylated pathway, which channels the glycolytic intermediate 3-phosphoglycerate (3PG) toward its biosynthesis ^21^. L-serine plays key roles in cell proliferation ^22^ and its biosynthetic rate-limiting enzyme, 3-Phosphoglycerate Dehydrogenase (PHGDH) is expressed by astrocytes, neural precursors and proliferating BrdU-positive cells, but not by neurons, in the DG of the adult mouse hippocampus ^21, 23.^ While most of L-serine is synthesized *de novo* in the brain, rise in its plasmatic concentration can result in a significant influx via the endothelial Slc38a5 transporter ^24^. L-serine contributes carbon to macromolecule biosynthesis and is converted to D-serine in the brain ^25^.

Through NMDAR signaling, D-serine supports the maturation, synaptic integration, and long-term survival of newly generated neurons in the adult hippocampus ^26, 27.^

Overall, these data suggest that a reduction in L-serine production during the course of AD may contribute to the deficits in AHN. Given that dietary supplementation can restore both L- and D-serine levels ^21^, we designed an experimental study to assess the effects of chronic L-serine supplementation on AHN in 3xTg-AD mice, a model that exhibits early metabolic alterations ^21^ and impaired AHN ^15^.

## 2. Materials and methods

### 2.1 Mouse models

All experimental procedures using animals were approved by a local ethics committee and submitted to the French Ministry of Education and Research (APAFIS#617-2015050516068503 v2 and APAFIS#30590). All mice were group-housed in individually ventilated cages with standard enrichment and maintained under conventional health status. Animals were kept at 22 ± 1 °C and 50% humidity on a 12 h light/dark cycle (lights on at 07:00) with food and water available ad libitum. We used 4-month-old homozygous female 3xTg-AD mice generated and maintained on a mixed B6;129 background (RRID:MMRRC_034830-JAX). 3xTg-AD mice express the mutated human genes *APP* Swe and *MAPT* P301L in the same locus under the control of the mouse Thy1.2 regulatory element as well as the mutated gene PS1M146V controlled by the cognate mouse *Psen1* locus ^28^. Age-matched control mice on the same genetic background were included as controls (B6129SF1/J; RRID:IMSR_JAX:101043).

### 2.2 Supplementation of L-serine

Four-month-old female 3xTg-AD (n = 24) and B6129SF1/J mice (n = 19) were fed either a standard diet containing 1% L-serine (w/w; Altromin 1310) or a 10% L-serine–enriched diet (w/w; Altromin 1324) for 3 or 8 months ^21^, see Figure 1 for the experimental design.

**FIGURE 1.**
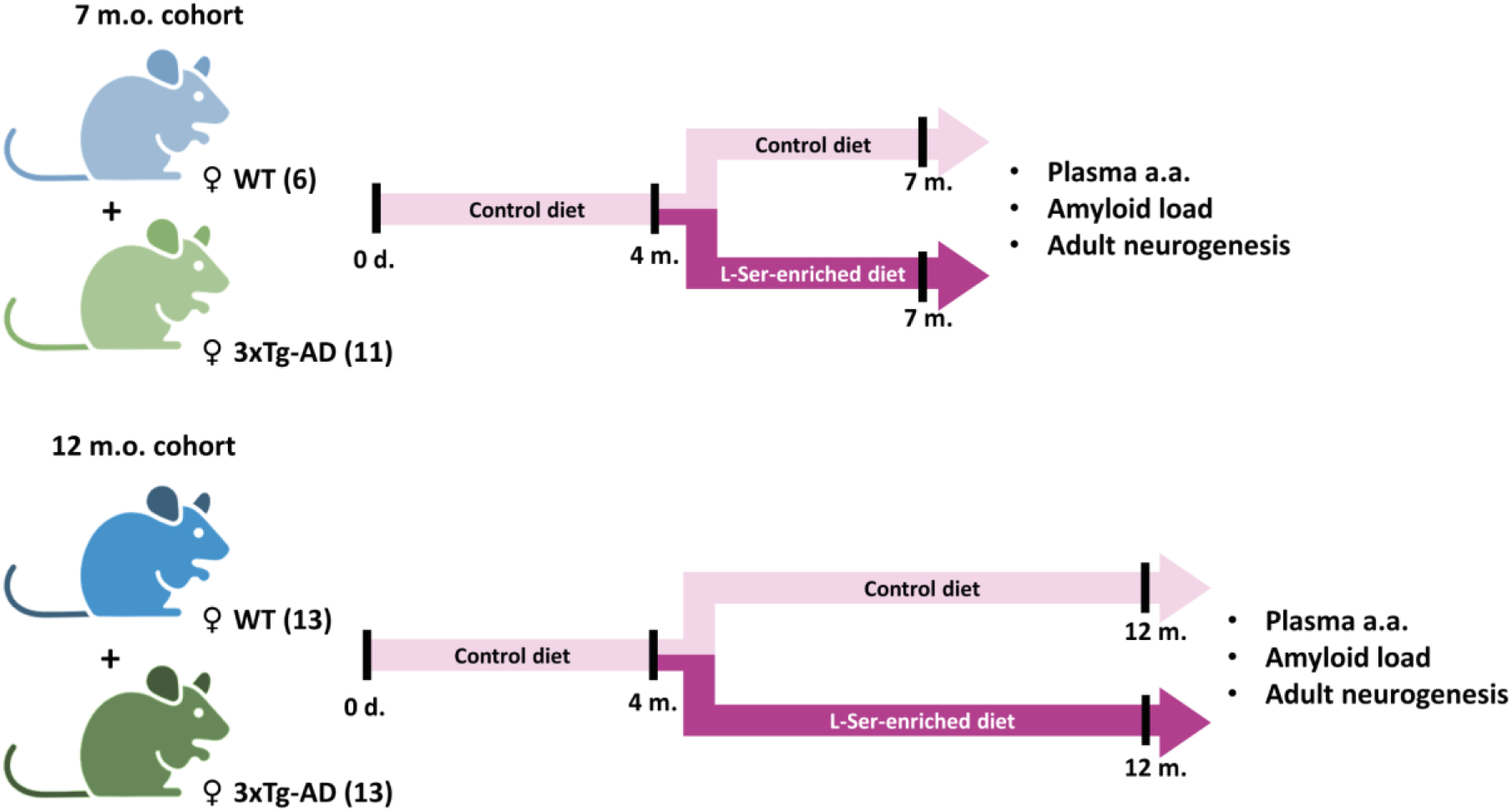
Experimental design showing the duration of dietary interventions and the number of animals per group (in parentheses). Mice were analyzed at either 7 or 12 months of age.

### 2.3 Determination of plasma amino acids

Mice were fasted for 2 hours starting at 9:00 a.m. and were sampled and transcardially perfused under anesthesia (see below). Blood was collected (50-300 µL) from the mandibular vein and stored in hemolysis tubes. Plasma was recovered in sterile tubes after centrifugation of blood samples at 2000 g and at 4°C for determination of L- and D-serine, glycine, glutamate, glutamine, leucine and methionine as previously described ^21^.

### 2.4 Immunohistochemistry of mouse brain for Aβ and PHGDH

Mice received a lethal dose of pentobarbital (400 mg/kg) by intraperitoneal injection and underwent intracardiac perfusion of 100 ml of cold 4% paraformaldehyde (PFA) solution in phosphate-buffered saline (PBS, 0.1 M, pH 7.4). The brains were cryoprotected in a 30% sucrose solution in PBS for 24 h. Sagittal brain sections (35 µm) were cut on a freezing microtome, sequentially collected into a series of six or seven 2 ml tubes (Eppendorf), and stored in a solution containing ethylene glycol, glycerol and 0.1 M PB at -20 °C until analysis. For Aß immunochemistry, brain sections were treated first with 70% formic acid for 2 minutes for antigen retrieval and immediately rinsed 3 × 10 minutes in 0.1 M PBS. They were then incubated in 0.3% H_2_O_2_ for 20 min and rinsed 3 × 10 minutes in PBS. After 30 minutes in blocking buffer (4.5% NGS, 0.2% Triton-X100 and PBS), sections were incubated for 24 h at 4°C with a biotinylated 4G8 antibody (mouse monoclonal; BioLegend, RRID:AB_2564651) diluted 1:1000 in a solution containing 3% NGS, 0.2% Triton-X100, and PBS. They were then incubated with the Vectastain Elite ABC Kit (PK-6100, Vector Laboratories) and revealed with the DAB-Nickel kit (SK-4100, Vector Laboratories). PHGDH immunostaining did not require pre-treatment with formic acid. Slices were incubated with primary (1:250, guinea pig, Frontier Institute, RRID:AB_2571654) and biotinylated secondary antibodies (Vector Laboratories), followed by Vectastain ABC and DAB-Nickel visualization.

### 2.5 Immunofluorescence staining of mouse brain

Cryopreserved brain sections were first rinsed 3 × 10 minutes in PBS and then incubated for 20 minutes at 80°C in citrate buffer for PCNA antigen retrieval. After 3 washes of 10 minutes in PBS, the sections were permeabilized and blocked for 1-2 hours in a solution of 0.2% Triton X-100 (Sigma) and 5% normal goat serum (NGS) or normal horse serum (NHS) in PBS. They were then incubated overnight at 4°C with the following primary antibodies diluted in a solution of 0.2% Triton-X100 and 3% NGS in PBS: proliferating cell nuclear antigen (PCNA) (1:500, rabbit, GeneTex, RRID:AB_1241163), doublecortin (DCX) (1:3000, guinea-pig, Merck Millipore, RRID:AB_1586992), PHGDH (1:250, guinea-pig, Frontier Institute, RRID:AB_2571654) and GFAP-Cy3 (1:500, mouse, Sigma, RRID:AB_476889). After three rinses in 0.2% Triton-X100 in PBS, brain sections were incubated for 3 h at RT with fluorescent secondary antibodies (1:1000, goat anti-rabbit Alexa Fluor 488, goat-anti guinea-pig Alexa Fluor 488 and goat anti-guinea pig Alexa Fluo 647, Invitrogen). After a brief rinse in 0.2% Triton-X100 in PBS, nuclei were stained with DAPI (0.1 µg/ml diluted in PBS) for 10 minutes. Lastly, sections were rinsed 3 more times in PBS and mounted on slides with FluorSave reagent (Calbiochem).

### 2.6 Human samples

Paraffin sections (hippocampus, parahippocampal gyrus and fusiform gyrus) of 2 control male brains (78 and 81 years-old) were provided by the GIE NeuroCEB biobank in Paris and were processed as previously described using a primary rabbit polyclonal PHGDH antibody (Frontier Institute, Hokkaido, Japan, RRID:AB_2571653) ^21^.

### 2.7 Image analysis

4G8 immunostained sections were scanned and digitized using an Axioscan Z1 slide scanner at x10 (Zeiss). Quantification of 4G8 staining was performed on a total of 11 sections per mouse brain. For each animal, all sections used for quantification were selected from the same part of the hippocampus, based on anatomical landmarks and extending approximately from +0.5 to +3.2 mm laterally from the midline. Images were imported with a 75% reduction in resolution and saved as “tif”. The area covered by the 4G8 staining was measured using thresholding algorithms in Fiji. The total area of 4G8 staining corresponds to the sum of the areas measured in all 11 sections.

Images (1024×1024) of fluorescent brain sections were acquired on a confocal microscope (Leica SP8) equipped with a 20X air objective (HC PL APO CS2 20x/0.75). Each line was averaged twice. An overlap of 20% was used to stitch image tiles. Image were analyzed on 3D reconstruction of z-stacks using the Imaris software (versions 9.8 and 9.9). Newborn immature neurons were defined as a cell displaying a DAPI-stained nucleus clearly associated with DCX immunostaining. PCNA-positive nuclei were automatically detected with the spot function of Imaris and then visually curated. Cell count in the DG was estimated by counting all cells in each section of a series covering the entire septotemporal extent of a hemisphere, then multiplying this total by the number of series obtained through sectioning.

### 2.8 Quantification and Statistical Analyses

No computational randomization methods were used to allocate animals between conditions. Statistical analyses were carried out using InVivoStat ^29^ and plots were created using Prism software (version 10). Residual normality was evaluated using normal probability plots and homogeneity of variance using predicted versus residual plots ^30^. Variables that deviated from these assumptions were systematically log10-transformed. This transformation was successful in normalizing the distribution of residuals and stabilizing the variance of the responses to make them homoscedastic. As a consequence, data were only analyzed using parametric tests. When several factors (treatment, genotype and age) were analyzed or/and when a factor presented more than two levels, overall effects were determined by analysis of variance (ANOVA). The overall effect of the different factors was not interpreted in the presence of significant interactions between them A gateway ANOVA strategy was not used. Pairwise comparisons were conducted using least significant difference (LSD) tests regardless of the ANOVA outcome. P-values were corrected for multiple comparisons using Hommel’s procedure. All statistical tests were two-tailed with significance set at α=0.05.

## 3. Results

### 3.1 PHGDH is expressed in NSCs of the DG in mice and Human

PHGDH catalyzes the first step of L-serine biosynthesis by oxidizing 3-phosphoglycerate (3PG) to 3-phosphohydroxypyruvate (3PHP). Using a specific antibody directed against PHGDH, we observed strong labeling in cells exhibiting typical astrocytic morphology, as well as in cells with somata located in the subgranular zone (SGZ) and displaying a radial process that penetrated the granule cell layer, a feature characteristic of NSCs in the DG of rodents (Figure 2A). PHGDH immunostaining was observed not only in the mouse brain but also in the human brain (Figure 2A). Confocal co-localization analyses further confirmed that PHGDH expression was predominantly associated with GFAP-positive NSCs in the mouse brain (Figure 2B). Such co-expression was observed in all 8 groups of mice. These observations confirm that the phosphorylated pathway is expressed by NSCs and not in post-mitotic neurons.

**FIGURE 2.**
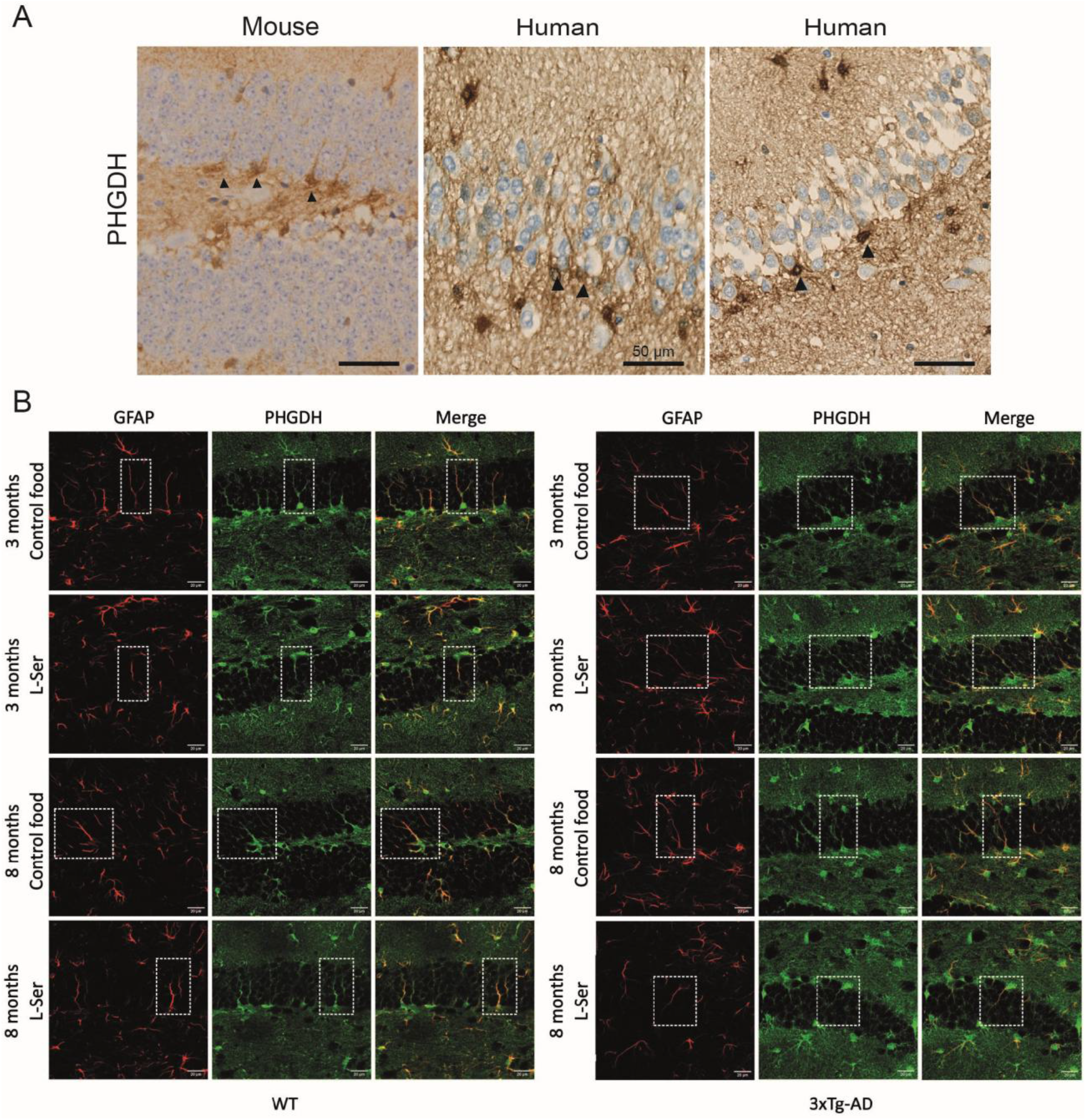
Expression of PHGDH in radial glia cells of the adult mouse and human DG. **A**. Immunostaining for PHGDH in the dentate gyrus (DG) of one adult mouse and two human brains. PHGDH is predominantly expressed in astrocytes and radial glia-like neural stem cells (NSCs), the latter identified by their radial processes and somata located in the subgranular zone (black arrows). **B**. Double immunofluorescent staining of brain sections from 7- and 12-month-old control and 3xTg-AD mice fed either a control or L-serine-enriched diet. Staining was performed using PHGDH (green) and the astrocyte/NSC marker GFAP (red).

### 3.2 L-Ser-enriched diet increases plasma L- and D-Serine levels

We then assessed the efficacy of an L-serine-enriched diet in selectively increasing plasma levels of L- and D-serine. All groups, except 3-month-old control mice, displayed higher plasma levels of L-serine when fed a diet enriched with 10% L-serine, with fold changes ranging from 2 to 3 (Figure 3). Unexpectedly, L-serine supplementation resulted in even higher increases in plasma D-serine, with levels reaching 9-to 22-fold those measured in mice fed the control diet (Figure 3). We also measured plasma levels of glycine, which is also synthesized from L-serine, and found no significant changes in response to either duration of L-serine treatment (Figure 3). This initial set of findings demonstrates the feasibility of specifically and chronically increasing the availability of plasma L- and D-serine via a peripheral delivery method, with the objective of promoting their transport into the brain.

**FIGURE 3.**
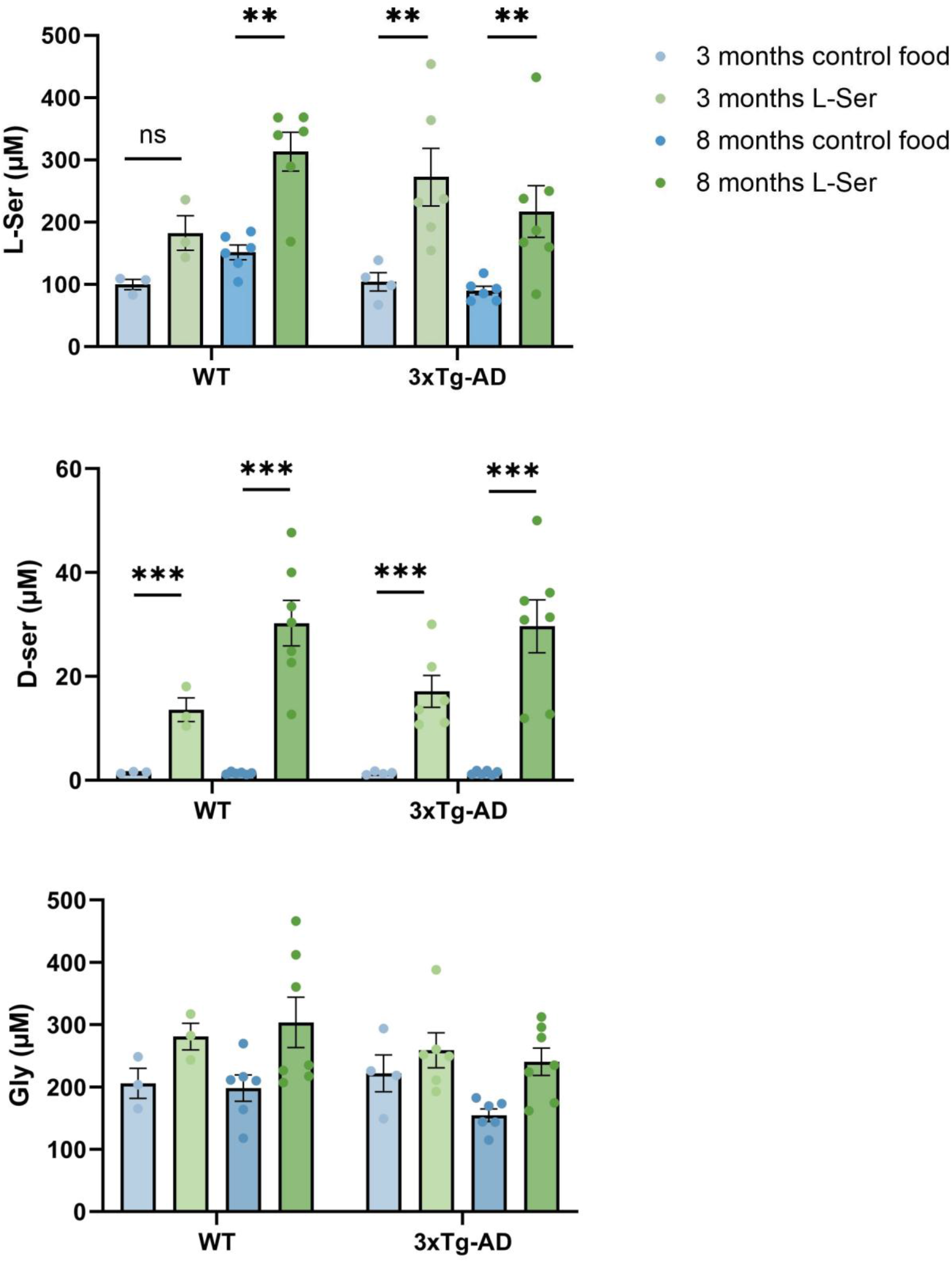
Plasma levels of L-serine, D-serine and glycine following 3- and 8-months of chronic L-serine dietary supplementation. **L-Ser**: Data were log10-transformed. Three-way ANOVA: genotype*duration*treatment: F(1,33)=0.36, p=0.5524; genotype*duration: F(1,33)=9.37, p=0.0044; genotype*treatment: F(1,33)=0.9, p=0.349; duration*treatment: F(1,33)=0, p=0.9726; genotype: F(1,33)=1.83, p=0.186; duration F(1,33)=1.6, p=0.215; treatment: F(1,33)=48.37, p<0.0001; A outlier was removed. Pairwise comparisons: LSD tests followed by Hommel’s correction for multiple comparisons. **D-Ser**: Data were log10-transformed. Three-way ANOVA: genotype*duration*treatment: F(1,34)=0.65, p=0.4242; genotype*duration: F(1,34)=0.04, p=0.8353; genotype*treatment: F(1,34)=0.16, p=0.6962; duration*treatment: F(1,34)=7.75, p=0.0087; genotype: F(1,34)=0.02, p=0.8954; duration F(1,34)=6.15, p=0.0183; treatment: F(1,34)=492.09, p< 0.0001; Pairwise comparisons: LSD tests followed by Hommel’s correction for multiple comparisons. **Gly**: Data were log10-transformed. Three-way ANOVA: genotype*duration*treatment: F(1,34)=0.28, p=0.5974; genotype*duration: F(1,34)=1.36, p= 0.2511; genotype*treatment: F(1,34)=0.18, p= 0.6756; duration*treatment: F(1,34)=1.11, p= 0.3005; genotype: F(1,34)=2, p= 0.1667; duration F(1,34)=1.71, p= 0.2004; treatment: F(1,34)=14.93, p=0.0005; Pairwise comparisons: LSD tests followed by Hommel’s correction for multiple comparisons. Mean with SEM.

### 3.3 L-Ser-enriched diet does not alter Aβ pathology in 3xTg-AD mice

We previously reported ^21^ that two months of an L-serine-enriched diet were sufficient to restore hippocampal L-serine levels in 3xTg-AD mice to values comparable to those of control mice, and to fully rescue the deficits in synaptic plasticity and spatial memory observed at 6 months of age. However, we did not address whether this dietary intervention impacted disease progression in terms of amyloid pathology. We therefore used the 4G8 β-amyloid monoclonal antibody to assess amyloid burden in two cohorts of 3xTg-AD mice that received the diet for 3 or 8 months, starting at 4 months of age. As already reported ^28^, 7-month-old female 3xTg-AD mice primarily exhibit intraneuronal 4G8 staining in several brain regions including the cortex, the hippocampus and the subiculum (Figure 4A). Notably, the DG was devoid of 4G8 immunostaining at both ages analyzed (Figure 4D). At 7 months, very few amyloid plaques were detected, whereas in 12-month-old female 3xTg-AD mice, plaque burden increased, particularly in the subiculum. Both small and large amyloid plaques were observed (Figure 4B), a pattern consistent with that seen in purely amyloidogenic mouse models such as 5xFAD ^31^ and TgAPParc ^32^. These findings align with post-mortem observations from AD patients and reinforce the notion that the subiculum is one of the earliest brain regions affected by the disease ^33^. Quantification of amyloid load revealed no significant effect of the L-serine-enriched diet compared to the control diet in either of the two 3xTg-AD cohorts (Figure 4C). We did not assess tau pathology, as it becomes significant only at later stages in this mouse model ^34^.

**FIGURE 4.**
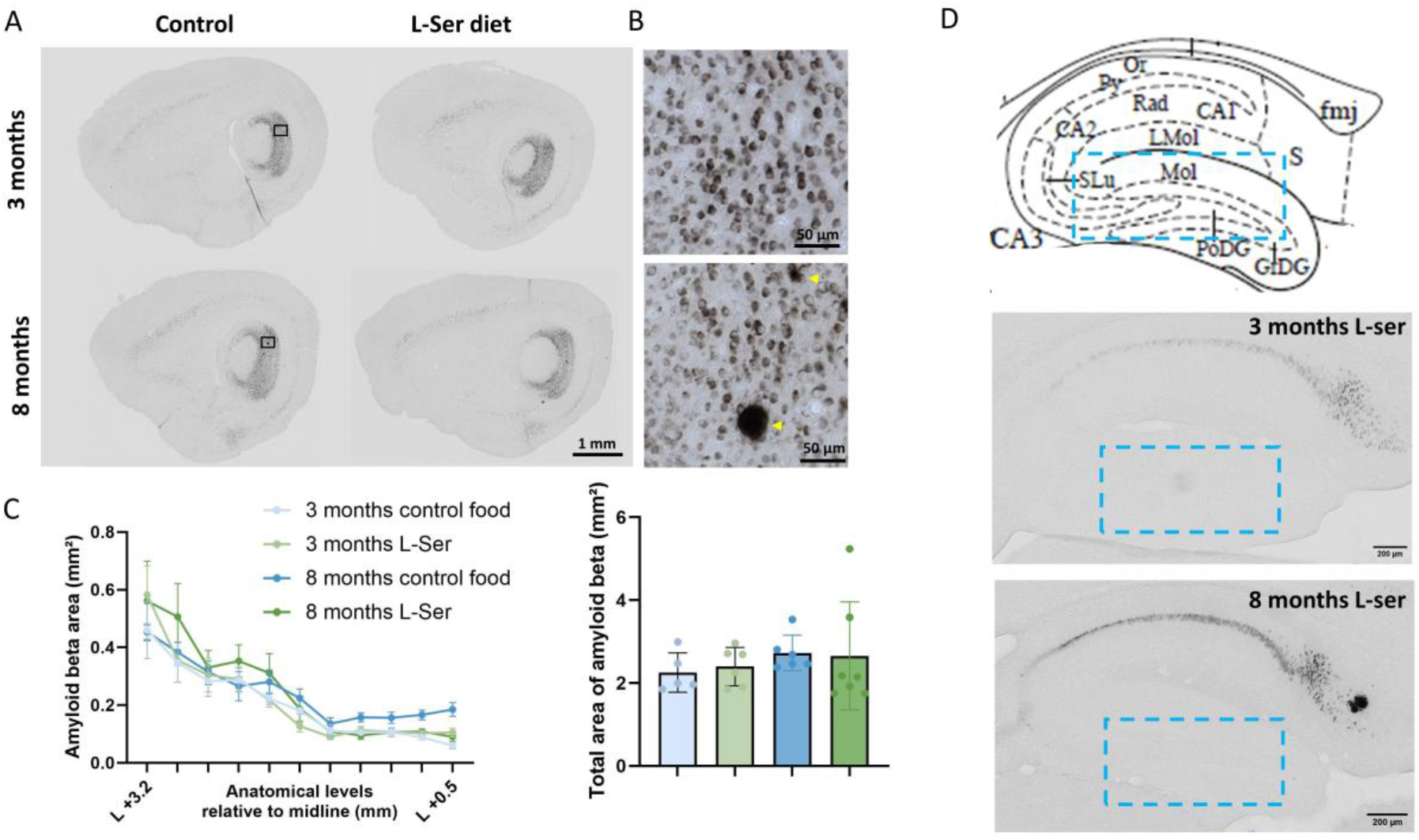
A Effect of L-serine dietary supplementation on amyloid load in 3xTg-AD mice. **A**. 4G8 staining in sagittal brain sections of 7 and 12-months-old 3xTg-AD mice that received 3 and 8 months of chronic control diet or L-serine dietary supplementation. **B**. Enlarged view of the CA1/Subiculum region (black boxes) showing that the immunostaining is mostly intraneuronal with few amyloid plaques (yellow arrowhead). **C**. Semi-quantitative analysis of 4G8-immunopositive staining performed in each of the 11 hippocampal sagittal sections (from +3.2 to +0.5mm lateral from the midline) shows no significant differences among the 4 groups of 3xTg-AD mice, as reflected in the histograms of total staining areas. **D**. The dentate Gyrus (DG, blue rectangle) is spared from amyloid deposits in 7 and 12 months-old 3xTg-AD mice that received 3 or 8 months of chronic L-serine dietary supplementation. Mean with SEM.

### 3.4 L-Ser-enriched diet partly rescues the adult neurogenesis deficit of 12-month-old 3xTg-AD mice

Next, we investigated the impact of two durations of L-serine supplementation (3 and 8 months) on the number of proliferating neural precursors and newborn neurons, identified by immunolabeling with PCNA and DCX, respectively. Accordingly, mice were studied at 7 and 12 months of age. We found that the number of DCX+ newborn neurons in control mice was significantly lower in 12-month-old mice compared with 7-month-old mice (354 ± 94 vs. 1772 ± 748 DCX+ cells, p < 0.0001), consistent with the established impact of aging on DG neurogenesis ^16^.

While no detectable reduction in immature DCX+ neurons was observed in 7-month-old 3xTg-AD mice, these animals exhibited a significant increase in proliferating neural precursors (Figure 5). In contrast, we observed a significant reduction in both DCX+ newborn neurons and proliferating neural precursors in the DG of 12-month-old 3xTg-AD mice compared to age-matched controls (Figure 6). Importantly, 8 months of an L-serine-enriched diet partially rescued both the number of proliferating neural precursors and the deficit in DCX+ immature neurons in 3xTg-AD mice, while having no significant effect on control mice (Figure 6).

**FIGURE 5.**
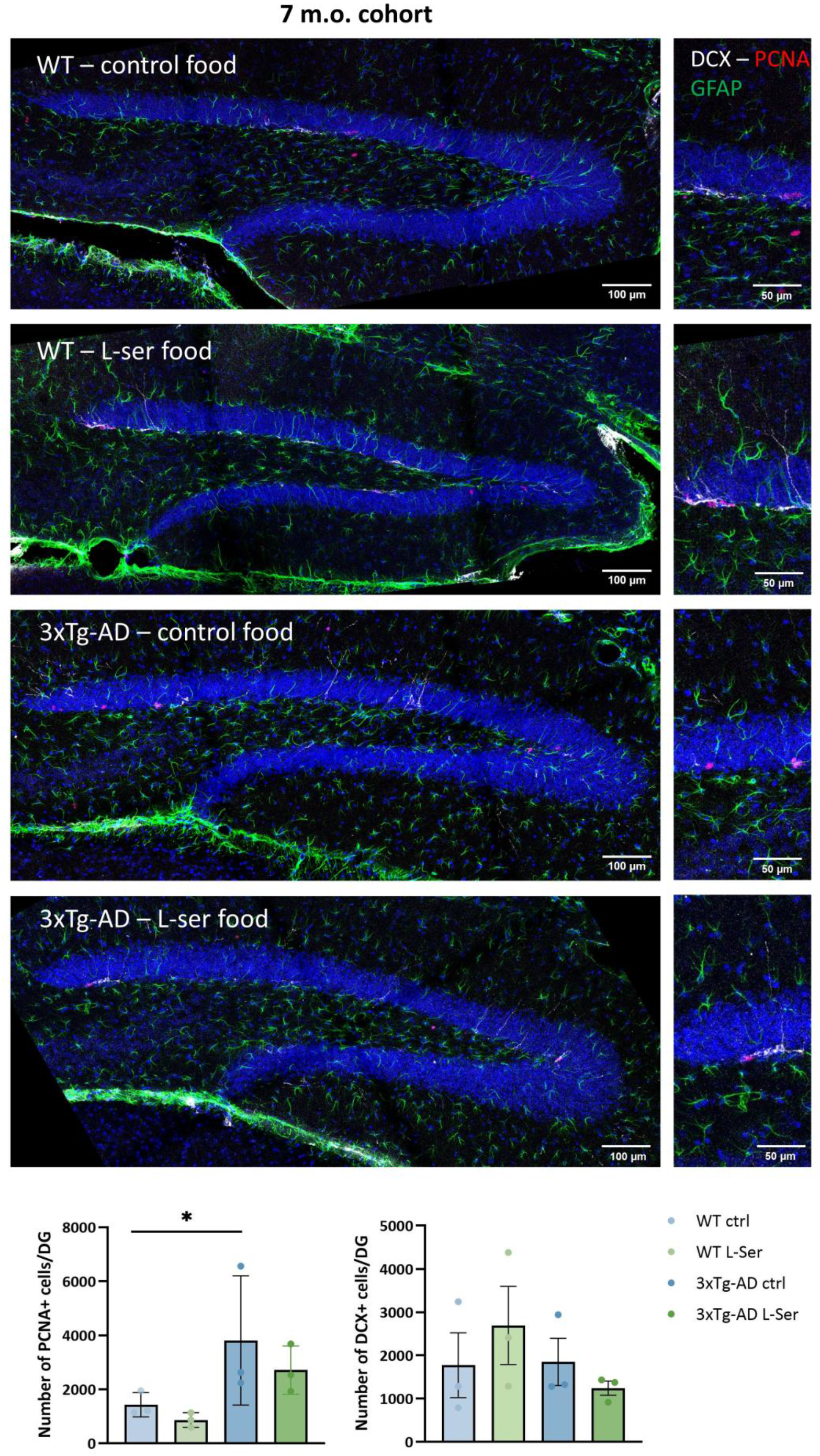
Effect of a 3-month L-serine diet on proliferating cells and immature neurons in the DG of WT and 3xTG-AD female mice. **A**. Representative images of hippocampal sections showing the DG from 7 month-old WT and 3xTg-AD mice fed either a control or a L-Ser-enriched diet for 3 months and immunostained for GFAP, DCX and PCNA to reveal astrocytes/NSCs, immature neurons and proliferating neural precursors, respectively. **B**. Quantification of proliferating precursors (left) and of immature neurons (right). Data were Log10-transformed prior to analysis. Three-way analysis of variance (ANOVA): 1) PCNA: Genotype * treatment * age, F(1,28) = 0.3, p = 0.5904; treatment * age, F(1,28) = 5.87, p = 0.0221; Genotype * age, F(1,28) = 35.24, p = < 0.0001; Genotype * treatment, F(1,28) = 2.31, p = 0.1399; Age, F(1,28) = 222.85, p = < 0.0001; Treatment, F(1,28) = 0.25, p = 0.621; Genotype F(1,28) = 3.4, p = 0.0758; 2) DCX: Genotype * treatment * age, F(1,28) = 4.77, p = 0.0375; treatment * age, F(1,28) = 0.85, p = 0.3635; Genotype * age, F(1,28) = 4.17, p = 0.0507; Genotype * treatment, F(1,28) = 0.01, p = 0.9387; Age, F(1,28) = 149.71, p = < 0.0001; Treatment, F(1,28) = 1.68, p = 0.2054; Genotype F(1,28) = 12.47, 0.0015; all pairwise comparisons: least significant difference (LSD) test followed by Hommel’s adjustment. Mean with SEM.

**FIGURE 6.**
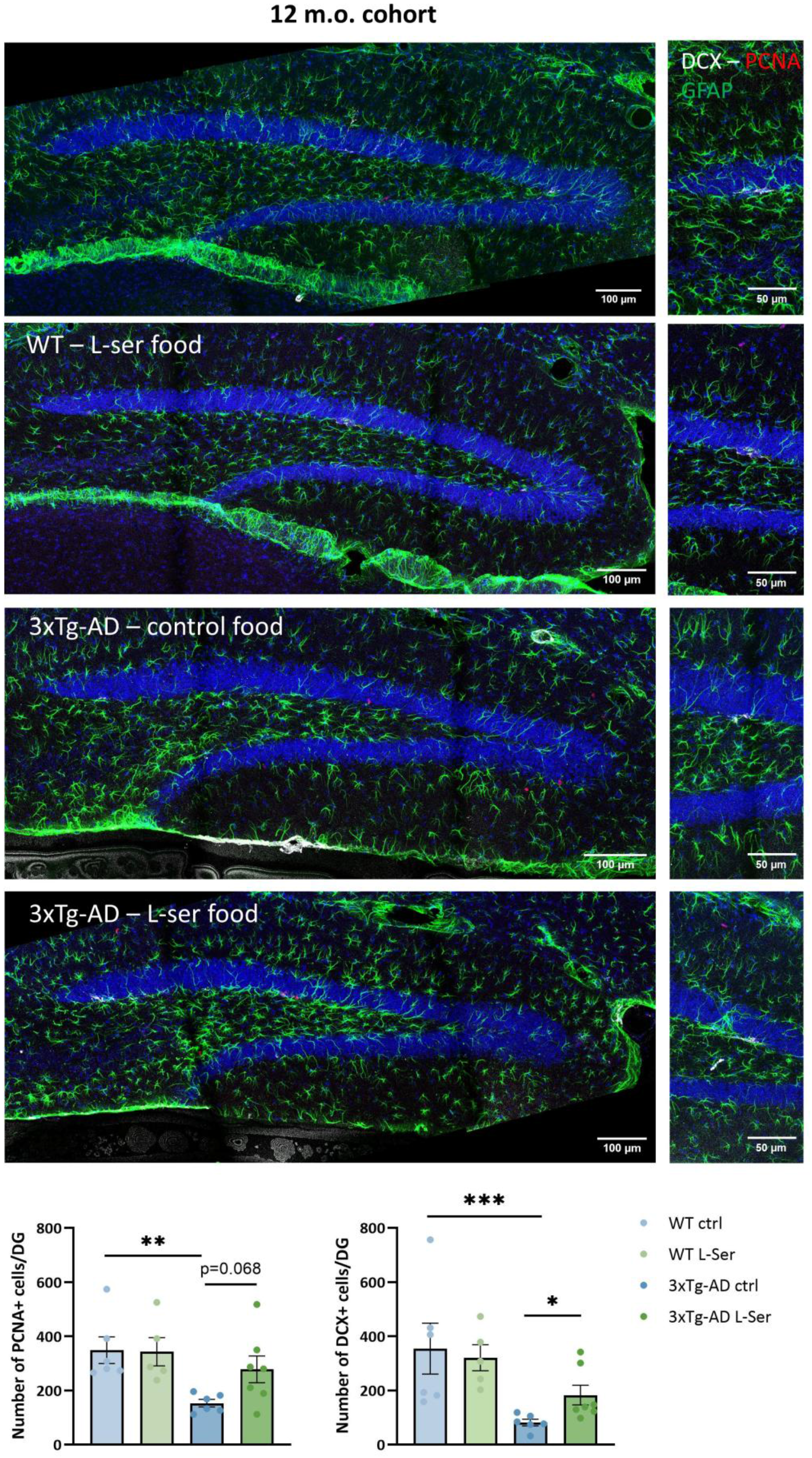
Effect of an 8-month L-serine diet on proliferating cells and immature neurons in the DG of WT and 3xTg-AD female mice. **A**. Representative images of hippocampal sections showing the DG from 12 month-old WT and 3xTg-AD mice fed either a control or a L-Ser-enriched diet for 8 months and immunostained for GFAP, DCX and PCNA to reveal astrocytes/NSCs, immature neurons and proliferating neural precursors, respectively. **B**. Quantification of proliferating precursors (left) and of immature neurons (right). Data were Log10-transformed prior to analysis. Three-way analysis of variance (ANOVA): 1) PCNA: Genotype * treatment * age, F(1,28) = 0.3, p = 0.5904; treatment * age, F(1,28) = 5.87, p = 0.0221; Genotype * age, F(1,28) = 35.24, p = < 0.0001; Genotype * treatment, F(1,28) = 2.31, p = 0.1399; Age, F(1,28) = 222.85, p = < 0.0001; Treatment, F(1,28) = 0.25, p = 0.621; Genotype F(1,28) = 3.4, p = 0.0758; 2) DCX: Genotype * treatment * age, F(1,28) = 4.77, p = 0.0375; treatment * age, F(1,28) = 0.85, p = 0.3635; Genotype * age, F(1,28) = 4.17, p = 0.0507; Genotype * treatment, F(1,28) = 0.01, p = 0.9387; Age, F(1,28) = 149.71, p = < 0.0001; Treatment, F(1,28) = 1.68, p = 0.2054; Genotype F(1,28) = 12.47, 0.0015; all pairwise comparisons: least significant difference (LSD) test followed by Hommel’s adjustment. Mean with SEM.

## 4. Discussion

Adult hippocampal neurogenesis is a relatively well-conserved process across mammalian species. It supports structural and functional plasticity underlying cognitive flexibility ^35^ and is influenced by environmental and behavioral factors that impact the number of newborn neurons ^16^. This suggests that the formation and integration of new neurons in the adult hippocampus remains plastic and potentially amenable to therapeutic intervention, even in the context of age-related decline and neurodegenerative conditions such as AD ^15^. In this study, we show that chronic dietary supplementation with L-serine - an amino acid essential for hippocampal neuron development ^36^ - partially mitigates the reduction in newborn neurons in the DG of 3xTg-AD mice.

The 3xTg-AD mouse model expresses *APP* Swe and *MAPT* P301L transgenes under the control of the Thy1.2 mini-gene promoter, which contains an estrogen response-like element resulting in higher transgene expression in females ^37^. Consistent with this, 3xTg-AD mice exhibit marked sexual dimorphism, with females developing amyloid plaques and neurofibrillary tangles earlier and more extensively than males ^34^. For this reason, only females were included in our study. Previous reports have consistently described impairments of DG neurogenesis in 3xTg-AD mice ^38-41^, although the reported timing of these deficits varies across studies. Importantly, AHN impairment in this mouse model precedes overt hippocampal pathology, including intraneuronal Aβ accumulation, extracellular plaque deposition, and Tau phosphorylation. Early disruptions of the SGZ neurogenic niche have been linked to intrinsic molecular alterations, including transcriptional changes in the Notch and Wnt pathways in neural stem/progenitor cells ^40^.

With the aim of chronically raising circulating levels of L- and D-serine and promoting their transport into the brain ^21^, mice were fed mice a diet enriched with 10% L-serine. Our mass spectrometry analyses demonstrate the high efficacy and selectivity of this dietary intervention, supporting the growing interest in L-serine supplementation as a therapeutic approach in various neurological disorders ^42^. In our previous work, we showed that this diet rescued hippocampal plasticity and spatial memory deficits in 7-month-old 3xTg-AD mice ^21^, but we did not assess its impact on amyloid pathology. Here, using 4G8 immunostaining, we show that chronic L-serine supplementation does not alter amyloid burden, indicating that the observed functional benefits occur independently of amyloid deposition. Consistent with prior reports, amyloid immunoreactivity was absent from the DG but prominent in the subiculum, where it increased with age ^34, 43^. Thus, prolonged elevation of circulating serine levels does not influence amyloid production or plaque accumulation in 3xTg-AD mice.

Our principal finding is that eight months of L-serine-enriched diet partially rescues the deficit in adult neurogenesis observed in 3xTg-AD mice. Unlike post-mitotic neurons, proliferating cells must generate sufficient biomass to support cell division, including nucleotides, lipids and amino acids. Accumulating evidence indicates that cellular metabolism is a key regulator of adult stem cell behavior *in vivo* ^44^. For example, lipid metabolism - encompassing both lipogenesis and fatty acid oxidation - is critical for NSC quiescence and proliferation ^45^. Among the amino acids influencing adult neurogenesis ^46^, L-serine occupies a central metabolic position, linking glycolysis-derived carbon to one-carbon metabolism, protein synthesis, lipid biosynthesis, and bioenergetic pathways ^42, 47.^ NSCs rely predominantly on glycolysis, whereas the transition toward neuronal differentiation is associated with increased oxidative phosphorylation, underscoring a tight coupling between metabolic state and neurogenic stage ^48, 49.^ Since most brain L-serine is synthesized locally from glycolysis-derived 3-phosphoglycerate ^50^, neural precursors and astrocytes, but not mature neurons, are likely contributors to its production. Transcriptomic analyses indeed indicate that, besides astrocytes, PHGDH is also expressed by quiescent NSCs within the adult DG ^51^, with PHGDH expression declining following NSC activation, in the course of lineage progression ^48^.

Surprisingly, despite our earlier observation that L-serine synthesis is already diminished in the hippocampus of 6-month-old 3xTg-AD mice (Le Douce et al., 2020), we did not detect any significant reduction in overall neurogenesis in 7-month-old 3xTg-AD mice. While the limited size of our cohort at that age calls for caution, the increased number of proliferating neural precursors that we still observed suggests the presence of a compensatory response, potentially aimed at maintaining a stable pool of DCX^+^ immature neurons in the face of reduced newborn neuron survival. Such a mechanism would ultimately lead to the exhaustion of the NSC pool, thus explaining the decrease in proliferating precursors and newborn neurons observed in older, 12-month-old 3xTg-AD mice. Such a mechanism is consistent with previous studies showing that premature or excessive recruitment of NSCs into the cell cycle ultimately results in stem cell exhaustion ^52-54^.

Taken together, our data support a model in which reduced L-serine availability, and more broadly altered anabolic metabolism in NSCs, contributes to impaired adult neurogenesis in 3xTg-AD mice. Our findings suggest that the neurogenic deficit observed in 12-month-old mice is primarily driven by reduced survival of newborn neurons rather than by impaired precursor proliferation, with early compensatory increases in NSC recruitment transiently masking this survival defect.

Several limitations of this study should be acknowledged. First, our experiments were conducted in a single AD mouse model, and it remains to be determined whether the beneficial effects of dietary L-serine supplementation generalize to other transgenic models. Comparisons with models lacking tau pathology may be particularly informative for dissecting the origins of serine deficiency. Second, our analysis relied on a limited set of classical neurogenesis markers and did not include BrdU pulse-chase experiments, which would have provided more direct insight into S-phase entry and newborn neuron survival. Finally, although behavioral assessment of pattern separation in aged and cognitively impaired mice is challenging, the inclusion of behavioral correlates would have strengthened the functional relevance of our findings.

## Acknowledgments

This work was supported by grants from Fondation Alzheimer (G.B.) and Fondation pour la Recherche Médicale (G.B.) as well as by INSERM (A.P., S.H.R.O.) and CNRS (A.P., S.H.R.O.). This work benefited from the support of various facilities granted by INSERM and the French government in the framework of the University of Bordeaux’s IdEx “Investments for the Future” program /GPR BRAIN_2030).

The authors thank Fanny Petit for her help with the histology, Delphine Gonzales, Sara Laumond and all the people of the Animal Facility and the Genotyping Facility of the NeuroCentre Magendie for mouse care and genotyping.

## References

1. Long JM and Holtzman DM. Alzheimer Disease: An Update on Pathobiology and Treatment Strategies. Cell 2019; 179: 312–339. DOI: 10.1016/j.cell.2019.09.001.

2. Spires-Jones TL and Hyman BT. The intersection of amyloid beta and tau at synapses in Alzheimer’s disease. Neuron 2014; 82: 756–771. DOI: 10.1016/j.neuron.2014.05.004.

3. Forner S, Baglietto-Vargas D, Martini AC, et al. Synaptic Impairment in Alzheimer’s Disease: A Dysregulated Symphony. Trends in Neurosciences 2017; 40: 347–357. DOI: 10.1016/j.tins.2017.04.002.

4. Braak H, Braak E and Bohl J. Staging of Alzheimer-related cortical destruction. Eur Neurol 1993; 33: 403–408. DOI: 10.1159/000116984.

5. Altman J and Das GD. Autoradiographic and histological evidence of postnatal hippocampal neurogenesis in rats. J Comp Neurol 1965; 124: 319–335. DOI: 10.1002/cne.901240303.

6. Seri B, Garcia-Verdugo JM, McEwen BS, et al. Astrocytes give rise to new neurons in the adult mammalian hippocampus. J Neurosci 2001; 21: 7153–7160. DOI: 10.1523/JNEUROSCI.21-18-07153.2001.

7. Toda T, Parylak SL, Linker SB, et al. The role of adult hippocampal neurogenesis in brain health and disease. Mol Psychiatry 2019; 24: 67–87. 20180420. DOI: 10.1038/s41380-018-0036-2.

8. Eriksson PS, Perfilieva E, Bjork-Eriksson T, et al. Neurogenesis in the adult human hippocampus. Nat Med 1998; 4: 1313–1317. DOI: 10.1038/3305.

9. Spalding KL, Bergmann O, Alkass K, et al. Dynamics of hippocampal neurogenesis in adult humans. Cell 2013; 153: 1219–1227. DOI: 10.1016/j.cell.2013.05.002.

10. Sorrells SF, Paredes MF, Cebrian-Silla A, et al. Human hippocampal neurogenesis drops sharply in children to undetectable levels in adults. Nature 2018; 555: 377–381. 20180307. DOI: 10.1038/nature25975.

11. Moreno-Jiménez EP, Flor-García M, Terreros-Roncal J, et al. Adult hippocampal neurogenesis is abundant in neurologically healthy subjects and drops sharply in patients with Alzheimer’s disease. Nature Medicine 2019; 25: 554–560. DOI: 10.1038/s41591-019-0375-9.

12. Salta E, Lazarov O, Fitzsimons CP, et al. Adult hippocampal neurogenesis in Alzheimer’s disease: A roadmap to clinical relevance. Cell Stem Cell 2023; 30: 120–136. DOI: 10.1016/j.stem.2023.01.002.

13. Tobin MK, Musaraca K, Disouky A, et al. Human Hippocampal Neurogenesis Persists in Aged Adults and Alzheimer’s Disease Patients. Cell Stem Cell 2019; 24: 974–982 e973. 20190523. DOI: 10.1016/j.stem.2019.05.003.

14. Zhou Y, Su Y, Li S, et al. Molecular landscapes of human hippocampal immature neurons across lifespan. Nature 2022; 607: 527–533. 20220706. DOI: 10.1038/s41586-022-04912-w.

15. Geigenmuller JN, Tari AR, Wisloff U, et al. The relationship between adult hippocampal neurogenesis and cognitive impairment in Alzheimer’s disease. Alzheimers Dement 2024; 20: 7369–7383. 20240821. DOI: 10.1002/alz.14179.

16. Babcock KR, Page JS, Fallon JR, et al. Adult Hippocampal Neurogenesis in Aging and Alzheimer’s Disease. Stem Cell Reports 2021; 16: 681–693. 20210225. DOI: 10.1016/j.stemcr.2021.01.019.

17. Rafalski VA and Brunet A. Energy metabolism in adult neural stem cell fate. Prog Neurobiol 2011; 93: 182–203. 20101105. DOI: 10.1016/j.pneurobio.2010.10.007.

18. Zheng X, Boyer L, Jin M, et al. Metabolic reprogramming during neuronal differentiation from aerobic glycolysis to neuronal oxidative phosphorylation. Elife 2016; 5 20160610. DOI: 10.7554/eLife.13374.

19. An Y, Varma VR, Varma S, et al. Evidence for brain glucose dysregulation in Alzheimer’s disease. Alzheimers & Dementia 2018; 14: 318–329. DOI: 10.1016/j.jalz.2017.09.011.

20. Batra R, Arnold M, Worheide MA, et al. The landscape of metabolic brain alterations in Alzheimer’s disease. Alzheimers Dement 2023; 19: 980–998. 20220713. DOI: 10.1002/alz.12714.

21. Le Douce J, Maugard M, Veran J, et al. Impairment of Glycolysis-Derived L-Serine Production in Astrocytes Contributes to Cognitive Deficits in Alzheimer’s Disease. Cell Metabolism 2020; 31: 503–517. DOI: 10.1016/j.cmet.2020.02.004.

22. Maugard M, Vigneron PA, Bolanos JP, et al. L-Serine links metabolism with neurotransmission. Progress in Neurobiology 2021; 197: 101896. DOI: ARTN 101896 10.1016/j.pneurobio.2020.101896.

23. Yamasaki M, Yamada K, Furuya S, et al. 3-Phosphoglycerate dehydrogenase, a key enzyme for l-serine biosynthesis, is preferentially expressed in the radial glia/astrocyte lineage and olfactory ensheathing glia in the mouse brain. J Neurosci 2001; 21: 7691–7704.

24. Radzishevsky I, Odeh M, Bodner O, et al. Impairment of serine transport across the blood-brain barrier by deletion of Slc38a5 causes developmental delay and motor dysfunction. Proc Natl Acad Sci U S A 2023; 120: e2302780120. 20231009. DOI: 10.1073/pnas.2302780120.

25. Ehmsen JT, Ma TM, Sason H, et al. D-serine in glia and neurons derives from 3-phosphoglycerate dehydrogenase. J Neurosci 2013; 33: 12464–12469. DOI: 10.1523/JNEUROSCI.4914-12.2013.

26. Sultan S, Li LY, Moss J, et al. Synaptic Integration of Adult-Born Hippocampal Neurons Is Locally Controlled by Astrocytes. Neuron 2015; 88: 957–972. DOI: 10.1016/j.neuron.2015.10.037.

27. Nacher J and McEwen BS. The role of N-methyl-D-asparate receptors in neurogenesis. Hippocampus 2006; 16: 267–270. DOI: 10.1002/hipo.20160.

28. Oddo S, Caccamo A, Shepherd JD, et al. Triple-transgenic model of Alzheimer’s disease with plaques and tangles: intracellular Abeta and synaptic dysfunction. Neuron 2003; 39: 409–421. 2003/08/05. DOI: S0896627303004343 [pii].

29. Clark RA, Shoaib M, Hewitt KN, et al. A comparison of InVivoStat with other statistical software packages for analysis of data generated from animal experiments. J Psychopharmacol 2012; 26: 1136–1142. 20111108. DOI: 10.1177/0269881111420313.

30. Bate ST and Clark RA. The Design and Statistical Analysis of Animal Experiments. Cambridge University Press, 2014.

31. Oakley H, Cole SL, Logan S, et al. Intraneuronal beta-amyloid aggregates, neurodegeneration, and neuron loss in transgenic mice with five familial Alzheimer’s disease mutations: potential factors in amyloid plaque formation. J Neurosci 2006; 26: 10129–10140. DOI: 10.1523/JNEUROSCI.1202-06.2006.

32. George S, Ronnback A, Gouras GK, et al. Lesion of the subiculum reduces the spread of amyloid beta pathology to interconnected brain regions in a mouse model of Alzheimer’s disease. Acta Neuropathol Commun 2014; 2: 17. 20140211. DOI: 10.1186/2051-5960-2-17.

33. Carlesimo GA, Piras F, Orfei MD, et al. Atrophy of presubiculum and subiculum is the earliest hippocampal anatomical marker of Alzheimer’s disease. Alzheimer’s & Dementia: Diagnosis, Assessment & Disease Monitoring 2015; 1: 24–32. DOI: 10.1016/j.dadm.2014.12.001.

34. Javonillo DI, Tran KM, Phan J, et al. Systematic Phenotyping and Characterization of the 3xTg-AD Mouse Model of Alzheimer’s Disease. Front Neurosci 2021; 15: 785276. 20220124. DOI: 10.3389/fnins.2021.785276.

35. Anacker C and Hen R. Adult hippocampal neurogenesis and cognitive flexibility - linking memory and mood. Nat Rev Neurosci 2017; 18: 335–346. 20170504. DOI: 10.1038/nrn.2017.45.

36. Mitoma J, Furuya S and Hirabayashi Y. A novel metabolic communication between neurons and astrocytes: non-essential amino acid L-serine released from astrocytes is essential for developing hippocampal neurons. Neurosci Res 1998; 30: 195–199.

37. Sadleir KR, Eimer WA, Cole SL, et al. Abeta reduction in BACE1 heterozygous null 5XFAD mice is associated with transgenic APP level. Mol Neurodegener 2015; 10: 1. 20150107. DOI: 10.1186/1750-1326-10-1.

38. Rodriguez JJ, Jones VC, Tabuchi M, et al. Impaired adult neurogenesis in the dentate gyrus of a triple transgenic mouse model of Alzheimer’s disease. PLoS One 2008; 3: e2935. 20080813. DOI: 10.1371/journal.pone.0002935.

39. Hamilton LK, Aumont A, Julien C, et al. Widespread deficits in adult neurogenesis precede plaque and tangle formation in the 3xTg mouse model of Alzheimer’s disease. Eur J Neurosci 2010; 32: 905–920. 20100819. DOI: 10.1111/j.1460-9568.2010.07379.x.

40. Liu Y, Bilen M, McNicoll MM, et al. Early postnatal defects in neurogenesis in the 3xTg mouse model of Alzheimer’s disease. Cell Death Dis 2023; 14: 138. 20230218. DOI: 10.1038/s41419-023-05650-1.

41. Wang JM, Singh C, Liu L, et al. Allopregnanolone reverses neurogenic and cognitive deficits in mouse model of Alzheimer’s disease. Proc Natl Acad Sci U S A 2010; 107: 6498–6503. 20100315. DOI: 10.1073/pnas.1001422107.

42. Handzlik MK and Metallo CM. Sources and Sinks of Serine in Nutrition, Health, and Disease. Annu Rev Nutr 2023; 43: 123–151. 20230612. DOI: 10.1146/annurev-nutr-061021-022648.

43. Lauritzen I, Pardossi-Piquard R, Bauer C, et al. The beta-secretase-derived C-terminal fragment of betaAPP, C99, but not Abeta, is a key contributor to early intraneuronal lesions in triple-transgenic mouse hippocampus. J Neurosci 2012; 32: 16243–11655a. DOI: 10.1523/JNEUROSCI.2775-12.2012.

44. Knobloch M and Jessberger S. Metabolism and neurogenesis. Curr Opin Neurobiol 2017; 42: 45–52. 20161201. DOI: 10.1016/j.conb.2016.11.006.

45. Knobloch M, Pilz GA, Ghesquiere B, et al. A Fatty Acid Oxidation-Dependent Metabolic Shift Regulates Adult Neural Stem Cell Activity. Cell Rep 2017; 20: 2144–2155. DOI: 10.1016/j.celrep.2017.08.029.

46. Guo Y, Luo X and Guo W. The impact of amino acid metabolism on adult neurogenesis. Biochem Soc Trans 2023; 51: 233–244. DOI: 10.1042/BST20220762.

47. Locasale JW. Serine, glycine and one-carbon units: Cancer metabolism in full circle. Nature Reviews Cancer 2013; 13: 572–583. DOI: 10.1038/nrc3557.

48. Shin J, Berg DA, Zhu Y, et al. Single-Cell RNA-Seq with Waterfall Reveals Molecular Cascades underlying Adult Neurogenesis. Cell Stem Cell 2015; 17: 360–372. 20150820. DOI: 10.1016/j.stem.2015.07.013.

49. Beckervordersandforth R, Ebert B, Schaffner I, et al. Role of Mitochondrial Metabolism in the Control of Early Lineage Progression and Aging Phenotypes in Adult Hippocampal Neurogenesis. Neuron 2017; 93: 1518. DOI: 10.1016/j.neuron.2017.03.008.

50. Yang JH, Wada A, Yoshida K, et al. Brain-specific Phgdh deletion reveals a pivotal role for L-serine biosynthesis in controlling the level of D-serine, an N-methyl-D-aspartate receptor co-agonist, in adult brain. J Biol Chem 2010; 285: 41380–41390. DOI: 10.1074/jbc.M110.187443.

51. Karpf J, Unichenko P, Chalmers N, et al. Dentate gyrus astrocytes exhibit layer-specific molecular, morphological and physiological features. Nat Neurosci 2022; 25: 1626–1638. 20221128. DOI: 10.1038/s41593-022-01192-5.

52. Ehm O, Goritz C, Covic M, et al. RBPJkappa-dependent signaling is essential for long-term maintenance of neural stem cells in the adult hippocampus. J Neurosci 2010; 30: 13794–13807. DOI: 10.1523/JNEUROSCI.1567-10.2010.

53. Mira H, Andreu Z, Suh H, et al. Signaling through BMPR-IA regulates quiescence and long-term activity of neural stem cells in the adult hippocampus. Cell Stem Cell 2010; 7: 78–89. DOI: 10.1016/j.stem.2010.04.016.

54. Urban N, van den Berg DL, Forget A, et al. Return to quiescence of mouse neural stem cells by degradation of a proactivation protein. Science 2016; 353: 292–295. DOI: 10.1126/science.aaf4802.

